# A survey of rare epigenetic variation in 23,116 human genomes identifies disease-relevant epivariations and novel CGG expansions

**DOI:** 10.1101/2020.03.25.007864

**Authors:** Paras Garg, Bharati Jadhav, Oscar L. Rodriguez, Nihir Patel, Alejandro Martin-Trujillo, Miten Jain, Sofie Metsu, Hugh Olsen, Benedict Paten, Beate Ritz, R. Frank Kooy, Jozef Gecz, Andrew J. Sharp

## Abstract

There is growing recognition that epivariations, most often recognized as promoter hypermethylation events that lead to gene silencing, are associated with a number of human diseases. However, little information exists on the prevalence and distribution of rare epigenetic variation in the human population. In order to address this, we performed a survey of methylation profiles from 23,116 individuals using the Illumina 450k array. Using a robust outlier approach, we identified 4,452 unique autosomal epivariations, including potentially inactivating promoter methylation events at 384 genes linked to human disease. For example, we observed promoter hypermethylation of *BRCA1* and *LDLR* at population frequencies of ~1 in 3,000 and ~1 in 6,000 respectively, suggesting that epivariations may underlie a fraction of human disease which would be missed by purely sequence-based approaches. Using expression data, we confirmed that many epivariations are associated with outlier gene expression. Analysis of SNV data and monozygous twin pairs suggests that approximately two thirds of epivariations segregate in the population secondary to underlying sequence mutations, while one third are likely sproradic events that occur post-zygotically. We identified 25 loci where rare hypermethylation coincided with the presence of an unstable CGG tandem repeat, and validated the presence of novel CGG expansions at several of these, identifying the molecular defect underlying most of the known folate-sensitive fragile sites in the genome. Our study provides a catalog of rare epigenetic changes in the human genome, gives insight into the underlying origins and consequences of epivariations, and identifies many novel hypermethylated CGG repeat expansions.

## INTRODUCTION

The main focus of the field of human genetics over the past few decades has been the investigation of sequence variation as a driver of human phenotypic variation. Projects such as the HapMap, 1000 Genomes, and the Exome Aggregation Consortium^1–5^ have provided deep surveys of genetic variation in both coding and non-coding regions, facilitating many novel insights into genotype-phenotype relationships in both common and rare diseases.

However, a number of recent studies have also demonstrated that rare epigenetic variation, termed epivariations or epimutations, can also underlie human disease. For example, between 5-15% of patients with hereditary nonpolyposis colorectal cancer who are negative for pathogenic coding variants present with constitutional *MLH1* [MIM: 120436] promoter methylation^6^. Similarly, allelic methylation of the *BRCA1* [MIM: 113705] promoter has been identified in several pedigrees with familial breast/ovarian cancer^7,8^, and inborn errors of vitamin B_12_ metabolism have been shown to result from an epivariation that silences *MMACHC*^9^ [MIM: 609831]. Other studies have shown a significant increase of *de novo* epivariations in individuals with congenital disorders compared to controls^10^, and provided evidence that epivariations contribute to the mutational spectra underlying autism and schizophrenia^11^.

Epivariations can be subdivided based on their apparent etiology^12^. Primary epivariations are thought to be caused by stochastic errors in the establishment or maintenance of the epigenome, such as certain types of imprinting anomalies^13^. In contrast, secondary epivariations occur as a result of an underlying change in local DNA sequence, and include mutations that disrupt regulatory elements^9,10,14^ and expansions of CpG-rich tandem repeats^15^. Large hypermethylated expansions of CGG repeats have been identified at a number of folate-sensitive fragile sites in the human genome^16^, including several that are associated with neurodevelopmental anomalies, such as the CGG expansions that occur at *FMR1* [MIM: 309550], *AFF2* [MIM: 300806], *DIP2B* [MIM: 611379] and *AFF3* [MIM: 601464]^17–20^.

Despite this growing evidence that epigenetic defects contribute to a wide variety of human diseases, currently little information exists on the prevalence and distribution of rare epigenetic variation in the human population. As a result, the potential contribution of epivariations to human disease is unclear. In order to address this, here we have analyzed data from >23,000 individuals that were originally generated for use in epigenome-wide association studies, representing the largest cohort of methylomes assembled to date. Utilizing a robust outlier analysis, we identified >4,000 epivariation loci that each span multiple CpGs in these samples, including several hundred that occur at the promoters of known Mendelian disease genes, thus implicating epivariations as a potentially causative factor in many human disorders. Using hundreds of monozygous (MZ) twin pairs and available SNP and expression data in thousands of samples, we investigated the causes and consequences of epivariations. Furthermore, by applying long-read sequencing, we validated the presence of novel CGG expansions as the cause of some epivariations, identifying the molecular defect underlying most of the known folate-sensitive fragile sites in the human genome.

Our study provides a catalog of rare epigenetic changes in the human genome, and identifies many novel hypermethylated CGG repeat expansions. We suggest that epivariations mark or represent a subset of pathogenic alleles at some disease loci, which would likely be missed by purely sequence-based approaches.

## MATERIALS AND METHODS

### Datasets

We accessed methylation data from a total of 24,985 individuals from 22 cohorts, listed in Table S1. Each cohort comprised DNA methylation profiles from at least 300 individuals generated using the Illumina 450k HumanMethylation BeadChip (450k array). Eighteen studies utilized DNA extracted from peripheral whole blood, while the remaining four studies utilized DNA extracted from newborn cord blood, dried neonatal blood spots, purified monocytes, or adipose tissue. Seventeen of the cohorts represented samples drawn from the general population without ascertainment for any specific condition, while five of the cohorts included some samples ascertained due to a diagnosis of ischemic stroke, asthma, Parkinson’s disease, facial clefts, or rheumatoid arthritis. Four of the cohorts were comprised partially or wholly of pairs of monozygous and dizygous twins. This study was approved by the Institutional Review Board of the Icahn School of Medicine under HS# 18-01169.

### Quality Control and Data Processing

Within each cohort, we performed a number of quality control steps to identify samples for exclusion, as follows: (i) We removed any sample with >1% of autosomal probes with detection p-value >0.01. (ii) We performed Principal Component Analysis (PCA) based on β-values of all probes located on chr1. Based on scatter plots of the first two principal components, we removed samples judged to be outliers. (iii) We utilized the array data to infer the likely sex of each sample, based on scatter plots of mean β-value of probes located on chrX versus the fraction of probes located on chrY with detection p>0.01. We compared these predictions against self-reported gender for each sample where available, and removed any samples with a potential sex mismatch. Furthermore, outlier samples, or samples with potential sex chromosome aneuploidies were also removed. (iv) We estimated the major cellular fractions comprising each blood sample directly from β-values using the method described by Houseman^21^. We then removed outlier samples, defined as those that showed cellular fractions either ≥99^th^ percentile +2%, or ≤1^st^ percentile −2% of any cell type. After these quality control steps, 23,173 samples remained for further processing and normalization, as described previously^10,11^. Briefly, raw signal intensities were subjected to color correction, background correction and quantile normalization using the Lumi package in R^22^, and the normalized intensities converted into β-values, which range between 0-1, representing the methylation ratio at each measured CpG. In order to correct for inherent differences in the distribution of β-values reported by Infinium I and Infinium II probes, we applied BMIQ^23^. Each cohort was normalized independently, and data for probes located on chrX in males were normalized separately from autosomal data.

### Identification of rare epigenetic variants

In order to identify rare epigenetic variants, also termed Differentially Methylated Regions (DMRs), we utilized a sliding window approach to compare individual methylation profiles of a single sample against all other samples from the same cohort. We chose this approach of testing for DMRs within each cohort in order to minimize batch effects that might result if we performed comparisons across different cohorts. We defined DMRs as regions of outlier methylation represented by multiple independent probes using the following parameters:

- Hypermethylated DMR: Any 1kb region with at least three or more probes with β-values ≥99.5^th^ percentile plus 0.15, and contains at least three consecutive probes with β-values ≥99.5^th^ percentile. In addition, we required that the minimum distance spanned by probes that were ≥99.5^th^ percentile was ≥100bp.
- Hypomethylated DMR: Any 1kb region with at least three or more probes with β-values ≤0.5^th^ percentile minus 0.15, and contains at least three consecutive probes with β-values ≤0.5^th^ percentile. In addition, we required that the minimum distance spanned by probes that were ≤0.5^th^ percentile was ≥100bp.

As the presence of an underlying homozygous deletion at a probe binding site can result in spurious β-values^11^, we removed any DMR call in which the carrier individual reported one or more probes within the DMR with failed detection p-value (p>0.01). Finally, we removed 57 samples which each reported an unusually high number (n>20) of autosomal DMRs, leaving a final cohort of 23,116 samples that were used in downstream analysis of autosomal loci. We performed manual curation of epivariation calls by visual inspection of plots, identifying 102 loci that showed clear technical effects, and that were removed.

For analysis of DMRs on the X chromosome, due to the confounder of X chromosome inactivation that can result in highly variable β-values at many X-linked loci in females, we only considered male samples in our analysis. Furthermore, to ensure statistical robustness for detecting outlier events, we only utilized chrX data from the ten cohorts that each contained at least 300 males after performing all QC steps (total n=8,027 males analyzed). Furthermore, due to hemizygosity for the X chromosome in males, which will result in stronger signals compared to heterozygous events on the autosomes, we increased thresholds for identifying DMRs on chrX to require three probes within a 1kb window with a β-value difference to ≥99.5^th^ percentile plus 0.4 for hypermethylated DMRs, and ≤0.5^th^ percentile minus 0.4 for hypomethylated DMRs. Before summarizing (Tables S2 and S3), overlapping DMRs identified in different individuals, but which showed methylation changes in the same direction, were merged.

We annotated DMRs using the following data sources: (i) Overlap with Refseq gene bodies and promoter regions (defined here as the region ±2kb of transcription start sites), (ii) Overlap with imprinted loci that exhibit significant parental bias in DNA methylation^24,25^. (iii) Overlap with repetitive elements identified by RepeatMasker and Tandem Repeats Finder (RepeatMasker and Simple Repeats tracks downloaded from the UCSC Genome Browser), (iv) OMIM disease genes based on overlap with Refseq gene promoters. All enrichment analyses were performed using a background list of 38,646 1kb windows on the 450k array that contained three or more probes, which overlap 68.8% of the 457,201 autosomal probes on the 450k array.

### Identification of candidate unstable tandem repeats

We utilized *hipSTR*^26^ to profile genome-wide variation of short tandem repeats (motif sizes 2-6bp) in a cohort of 600 individuals who had undergone whole genome sequencing using Illumina 150bp paired-end reads, representing the parents of individuals with congenital heart defects (dbGaP Study Accession: phs001138.v3.p2).

### Analysis of monozygotic twins

For concordance analysis of epivariations found in MZ twins, we generated β-value plots of each epivariation identified in any MZ twin, and used these to manually categorize each locus as fully concordant, partially concordant, or discordant within each MZ twin pair.

### Analysis of gene expression data

Four of the cohorts utilized in this study had available gene expression data, as follows:

1. BIOS study: We downloaded gene-level RNAseq read counts for 3,560 samples made using HTSeq (European Genome-phenome Archive dataset EGAD00010001420)^27^. Read counts were normalized using DESeq2^28^. We only considered autosomal genes with mean expression value in the top half of all genes assayed.
2. MuTHER study: We used normalized expression values for 825 samples with expression in subcutaneous fat generated using the Illumina HumanHT-12 v3.0 Expression BeadChip (Array Express dataset E-TABM-1140). Probe sequences were mapped using BWA, and only uniquely aligned probes were retained. We removed any probe that overlapped with SNVs identified by the 1000 Genomes Project that had Minor Allele Frequency (MAF)>0.01 in European populations, and only considered autosomal genes with mean expression value in the top half of all genes assayed.
3. MESA study: We used normalized expression values for 1,202 samples generated using the Illumina HumanHT-12 v4.0 expression beadchip (GEO accession GSE56045). We removed any expression value with detection p>0.01, removed probes with more than 10% missing values, and only considered autosomal genes with mean expression value in the top half of all genes assayed.
4. Framingham Heart Study: We used normalized expression values for 2,198 samples generated using the Affymetrix GeneChip Human Exon 1.0 ST Array (dbGaP Study Accession: phs000363.v5.p7). We only considered autosomal genes with mean expression value in the top half of all genes assayed.

In each cohort, we linked epivariations to corresponding expression data based on the overlap of epivariations with Refseq gene promoters (as defined above), retaining only those genes that showed a unique mapping position with a single gene promoter. Normalized gene expression values were converted to both z-scores and ranks, and we compared expression data for samples carrying hypomethylated epivariations or hypermethylated epivariations against the entire population. P-values were generated by randomly permuting expression values 10,000 times among samples, and comparing the mean gene expression of these permuted values with the observed means of genes associated with epivariations.

### *Cis*-association analysis of epivariations with SNVs

We used available SNV array data from 933 samples from the WHI cohort genotyped with the Illumina Multi-Ethnic Genotyping Array for whom methylation data were also available.

We performed pre-imputation quality control on the raw SNV array data which included removing multi-allelic sites, indels, resolving strand inconsistencies, and converting coordinates from hg18 to hg19, where applicable, using PLINK (versions 1.07 and 2b3.43)^29,30^, vcftools (version 0.1.15)^31^ and Beagle utilities. We performed imputation and phasing in each of the datasets separately using Beagle 4.0^32^ and the 1000 Genomes Project (1KGP) Phase3 reference panel downloaded from the Beagle website. For efficiency, genotype data from each chromosome was divided into segments of 5,000 SNVs for imputation, processed separately, and subsequently merged together for downstream analysis. We performed quality control on imputed and phased genotypes, removing SNVs with imputed R^2^<0.95, Hardy-Weinberg equilibrium p<10^−4^, and multiallelic sites.

We selected 97 epivariations that were present in ≥2 or more individuals in the WHI cohort and performed a χ-square test using SNVs located within ±1Mb around each epivariation, comparing allele frequencies between epivariation carriers and all other samples who did not carry that epivariation. We considered SNVs as significantly associated at 1% FDR.

### Validation of repeat expansions using long read sequencing

Pacific Biosciences long insert libraries with the addition of barcodes were prepared for samples with epivariations at *ABCD3* and *PCMTD2*, the two samples mixed at equimolar amounts, and sequenced on a single 8M SMRT cell with the Pacific Biosciences Sequel II system. Mean coverage was 12.5x and 9.1x, mean polymerase read lengths were 35.8kb and 34.2kb, and mean subread lengths were 10.7kb and 10.0kb for samples with epivariations of *PCMTD2* and *ABCD3*, respectively. Subreads were aligned to the hg19 human reference genome using pbmm2 v1.0.0^33^ with default parameters. Subreads were extracted from hg19 coordinates chr1:94,883,969-94,884,008 and chr20:62,887,069-62,887,108, and the number of CGG motifs were detected using the TR-specific dynamic programming algorithm *PacmonSTR*^34^ from the extracted subreads. We sequenced samples with epivariations at *LINGO3* and *FZD6* using Oxford Nanopore Technology, generating mean coverage of 3x and 27x, respectively. Reads were mapped to the hg19 human reference genome using minimap2 (version 2.7)^33^, and bam files for samples sequenced in multiple runs were merged, sorted and indexed using samtools (version 1.7)^35^. To estimate methylation levels on normal and expanded CGG repeat alleles separately, we first separated reads in each sample based on the presence or absence of a CGG expansion. Using nanopolish (version 0.10.2)^36^, we created index files to link reads with their signal level data in FAST5 files, followed by estimation of DNA methylation status at each CpG located within 2kb of CGG TRs, requiring a minimum log likelihood ratio ≥2.5 at each site.

### Southern blot, repeat-Primed PCR, methylation and expression analysis in a carrier of FRA22A

A Southern blot was created by digesting 8μg DNA extracted from peripheral blood, using restriction enzymes *Hind*III and *Xba*I. The digested DNA was then separated by electrophoresis on a 0.7% agarose gel, and after denaturation and neutralization, transferred to Hybond N+ membranes. Hybridization was performed at 65°C using a specific probe generated by PCR (forward primer GCTGGAGAGGGAGGGAAGG and reverse primer ATAGAAACGAAGGCAAAGGAGACC).

Repeat-primed PCR was performed to interrogate the number of CGG repeats in the *CSNK1E* gene with the Asuragen CGG Repeat Primed PCR system designed for detection of Fragile X expanded alleles. Samples were PCR-amplified using 2μl of DNA sample (20 ng/μl), 11.45μl of GC-rich AMP buffer, 0.25μl of FAM-labeled *CSNK1E* forward primer F1 (AGGCTGGGGAACTGCGTCT) or FAM-labeled *CSNK1E* forward primer F2 (GAGAGCCCAGAGCCAGAGC), 0.25μl of *CSNK1E* reverse primer R3 (CAAAAACAAAGAGGCTGAGGGAG), 0.5μl of CGG primer (TACGCATCCCAGTTTGAGACGGCCGCCGCCGCCGCC), 0.5μl of nuclease-free water, and 0.05μl of GC-rich polymerase mix from Asuragen Inc. (Austin, TX). Samples were amplified with an initial heat denaturation step of 95°C for 5 min, followed by 10 cycles of 97°C for 35 sec, 62°C for 35 sec, and 68°C for 4 min, and then 20 cycles of 97°C for 35 sec, 62°C for 35 sec, and 68°C for 4 min, with a 20 sec auto extension at each cycle. The final extension step was 72°C for 10 min. After PCR, 2μl of the PCR product was added to a mix with 11μl formamide and 2 μl Rox 1000 size standard (Abbott, Abbott Park, IL). After a brief denaturation step, samples were analyzed using an ABI Prism 3130 Genetic Analyzer (Applied Biosystems, Foster City, CA).

DNA methylation analysis was performed using bisulphite treatment with the Epitect bisulfite kit (Qiagen, Venlo, Netherlands). Primers specific for the methylated bisulphite converted DNA (GAGGAGGAGGGGGTTTGTTAT and AAATCAATAACCTAATAACCACACAC) were designed using Methyl Primer Express (Applied Biosystems, Foster City, CA, USA). After PCR amplification, the CGG surrounding area was sequenced using the forward primer on an ABI Prism 3130 (Applied Biosystems, Foster City, CA, USA). We performed pyrosequencing to quantify the methylation using the *CSNK1E*_001 PyroMark CpG assay and analyzed the results on a PyroMark Q24 (Qiagen, Venlo, Netherlands). Methylation threshold values used were 10%.

Quantitative RT-PCR analysis was used to assess expression levels of the *CSNK1E* gene. After homogenizing cultured lymphoblastoid cells from the FRA22A-expressing individual in triplicate, and from nine control individuals, total cellular RNA was isolated using Trizol (Invitrogen, Carlsbad, CA) according to the manufacturer’s instructions, with RNase-free DNase treatment (Ambion, Austin, TX). Subsequently, cDNA was reverse transcribed from total patient and control RNA samples using random hexamers primers from the SuperScript™ III First-Strand Synthesis System for RT-PCR kit (Invitrogen, Carlsbad, CA) according to manufacturer’s guidelines. Genomic contamination of the cDNA was checked with 2 control primers (ATAGTCACCCCATTCAAACTCAAG and ATTCATAGCAGCAGCATTTGTTTTA), spanning a large intron. First strand cDNA was diluted in TE buffer to a concentration of 20 ng/μl. Primers were designed to span the exon-exon junction between protein coding exons 6 and 7 of *CSNK1E* (TCAGCGAGAAGAAGATGTCAAC and GTAGGTAAGAGTAGTCGGGC), and mRNA expression assayed with a two-step real-time quantitative PCR assay with qPCR MasterMix Plus w/o UNG with SYBR^®^ Green I No Rox (Eurogentec S.A, Seraing, Belgium) using a Lightcycler^®^ 480 Instrument (Roche Applied Science, Basel, Switzerland). Cycling conditions were as follows: An UNG step of 2 min 50°C, 10 min 95°C and 45 cycles at 95°C for 15 sec and 60° C for 1 min. Subsequently, specificity of the amplification was checked using a melting curve analysis by rapid heating to 97°C to denature the DNA (11°C/s), followed by cooling to 65°C (0.4°C/s). The protocol was terminated with a cooling step of 10 sec at 40°C. All samples were assayed in triplicate. Expression of *CSNK1E* was normalized against the geometric mean of three stably expressed reference genes (*B2M, GAPDH* and *YWHAZ*), and a Mann Whitney U test was used to assess statistical significance.

## RESULTS

Using a sliding window approach to identify regions containing ≥3 CpGs with strong outlier methylation levels (see Methods), we identified 13,879 curated autosomal epivariations in 7,653 individuals, and 26 chrX epivariations in 26 males. Overall, 33.1% of the 23,116 samples tested carried one or more epivariations, corresponding to 4,452 unique autosomal loci, and 18 unique chr X loci (Tables S2 and S3). We observed an ~2.3-fold excess of hypermethylated compared to hypomethylated epivariations: of the autosomal loci, 3,095 epivariations were gains of methylation, 1,329 epivariations were losses, while 28 autosomal epivariations were bidirectional, exhibiting either hyper- or hypomethylation in different samples.

Given the size of our cohort, we were able to estimate the population frequency of each epivariation (Tables S2 and S3), including several that have been described previously and/or are associated with disease. For example, the second most frequent epivariation we observed was hypermethylation of the promoter region of *FRA10AC1* [MIM: 608866], with a population frequency of ~1 per 325 individuals. This epivariation is known to be caused by expansion of an underlying CGG repeat which causes silencing of *FRA10AC1*, and in heterozygous form is thought to be a benign variant^37^. Similarly, we observed gains of methylation at *DIP2B* with a frequency of ~1 per 1,050 samples, and *XYLT1* [MIM: 608124] in ~1 per 2,100 samples. Both of these events are also caused by underlying expansions of CGG repeats and have been associated with intellectual disability^19^ and recessive Desbuquois dysplasia 2, respectively^15^. Other known epivariations we observed include promoter methylation of *MMACHC*, which can cause recessive inborn errors of vitamin B_12_ metabolism^9^ and which we observed at a population frequency of ~1 in 950. The frequency distribution of hyper- and hypomethylated autosomal epivariations is shown in Figure S1.

2,723 (61.2%) of 4,452 epivariations overlapped broad gene promoter regions (±2kb of transcription start site), including 499 (402 hypermethylated, 91 hypomethylated, and 6 bidirectional epivariations) that overlapped promoter regions of OMIM disease genes. We observed evidence suggestive of purifying selection operating on promoter-associated epivariations (Figure 1). Using pLI scores generated by the Exome Aggregation Consortium (ExAC)^5^, hypermethylated promoter epivariations were biased away from the promoters of genes under selective constraint (permutation p<10^−7^). Similarly, hypomethylated epivariations also showed bias away from constrained genes (permutation p=1.6×10^−3^), but to a lesser degree than hypermethylated epivariations. We also observed a weak but significant inverse relationship between the population frequency of hypermethylated promoter epivariations and selective constraint of the associated genes (Pearson r=-0.11, p=1.8×10^−6^) (Figure 1b).

**Figure 1.**
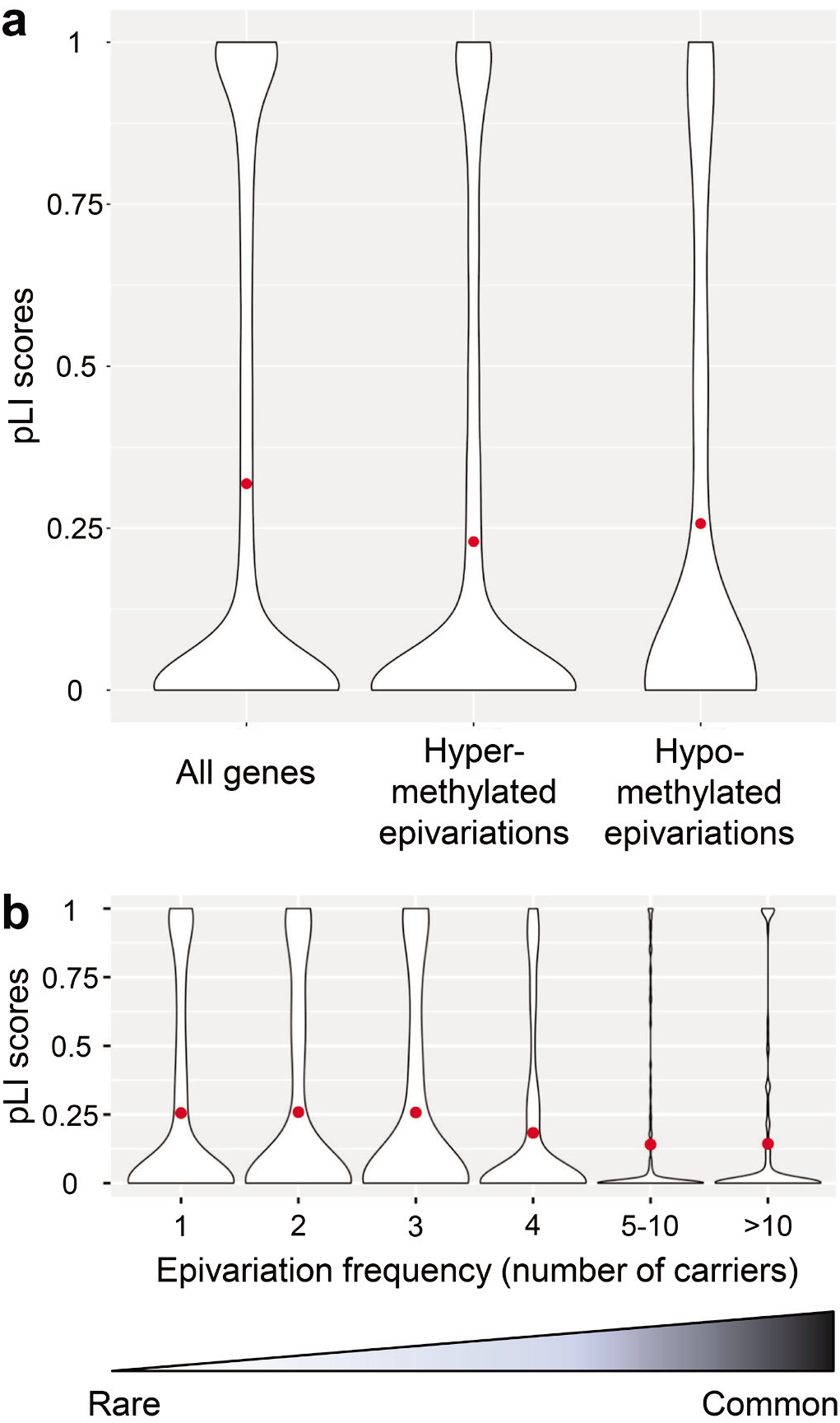
Evidence of purifying selection operating on promoter epivariations. **(A)** Using pLI scores generated by the ExAC^5^, we observed that hypermethylated promoter epivariations were preferentially associated with genes under selective constraint (permutation p<10^−7^). Similarly, hypomethylated epivariations were also biased away from genes with high pLI scores, but to a lesser degree (permutation p=1.6×10^−3^). **(B)** We observed an inverse relationship between the population frequency of hypermethylated promoter epivariations and selective constraint of the associated gene (Pearson p=1.8×10^−6^). These distributions are consistent with many promoter epivariations undergoing purifying selection which reduce their frequency in the population.

### Epivariations are frequently associated with *cis*-linked changes in gene expression

To determine the functional effect of epivariations on local gene expression, we analyzed available gene expression data in four different cohorts, comprising a total of 7,786 samples, analyzed with three different expression platforms. Focusing on epivariations that occurred at the promoter regions of genes, we observed significantly altered gene expression levels associated with epivariations in every cohort (Figure 2). Consistent with the known repressive effects of promoter methylation^38^, promoter hypomethylation was associated with increased expression in all four cohorts tested, while promoter hypermethylation was associated with reduced repression (Tables S4-S7).

**Figure 2.**
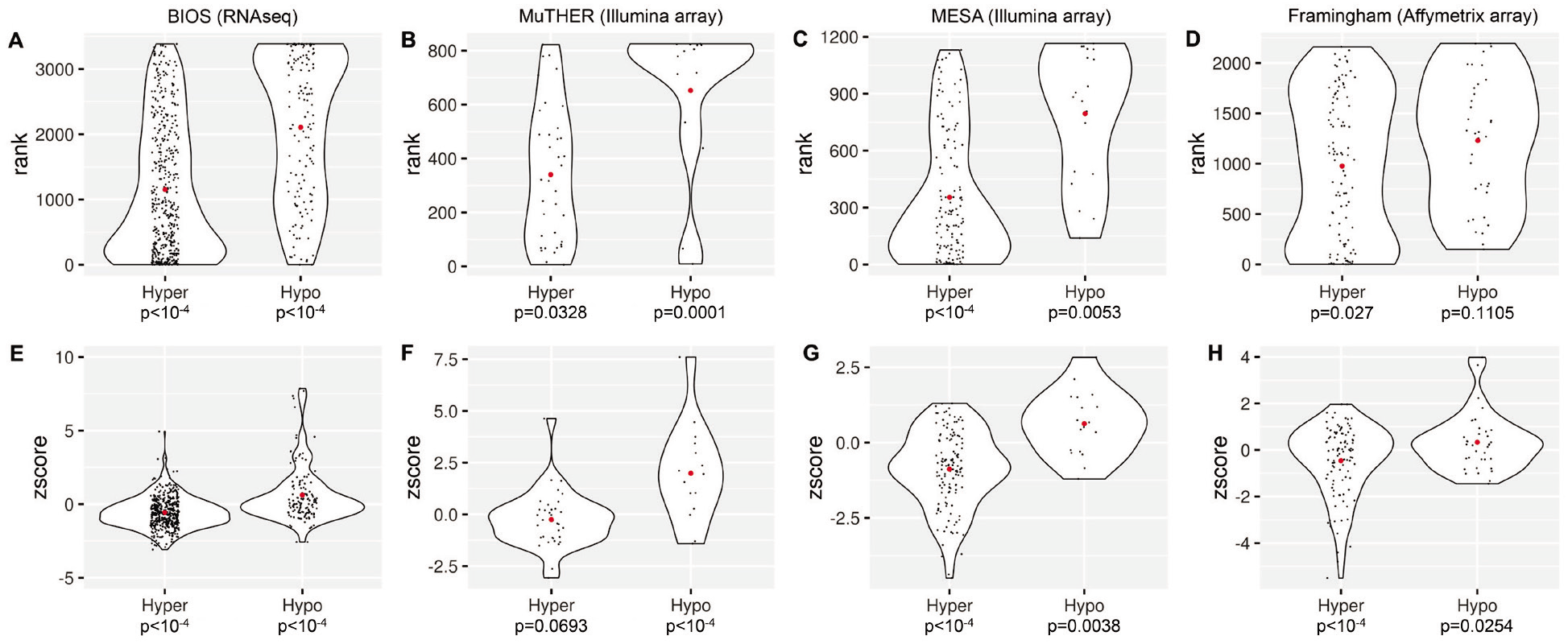
Promoter epivariations are associated with altered gene expression. **(A-D)** Rank distributions of expression for genes with promoter epivariations, and **(E-H)** z-scores of expression for genes with promoter epivariations. Using available gene expression data in four different cohorts, we observed that epivariations located at promoter regions were associated with altered gene expression levels. Consistent with the known repressive effects of promoter DNA methylation, promoter hypermethylation was associated with transcriptional repression, and promoter hypomethylation with increased expression. In each violin plot, the red dot shows the mean expression value of each distribution, with individual genes shown as black dots. Above each plot is shown the cohort name and expression platform.

We also performed similar tests of the effect on expression of epivariations that either overlapped gene bodies (excluding promoter regions), and for the effects of intergenic epivariations on the closest gene. Using RNAseq data from the BIOS cohort, we observed small but nominally significant effects of gene body methylation, which showed a weak positive correlation with expression (Figure S2). However, using closest gene annotations, we observed no significant associations.

### Epivariations and known disease genes

To gain insight into the potential contribution of epivariations to human disease, we utilized OMIM disease gene annotations, identifying 384 autosomal OMIM genes with hypermethylated epivariations at their promoter regions that may result in allelic silencing (Table S2). This includes seven of 59 genes in which pathogenic mutations are considered medically actionable by the American College of Medical Genetics^39^. For example, we detected seven individuals with promoter methylation of *BRCA1*, which has previously been reported in pedigrees with hereditary breast/ovarian cancer who lack pathogenic coding mutations in *BRCA1*^7,8^, and four individuals with promoter methylation of *LDLR* [MIM: 606945], haploinsufficiency of which is associated with familial hypercholesterolemia (Figure 3).

**Figure 3.**
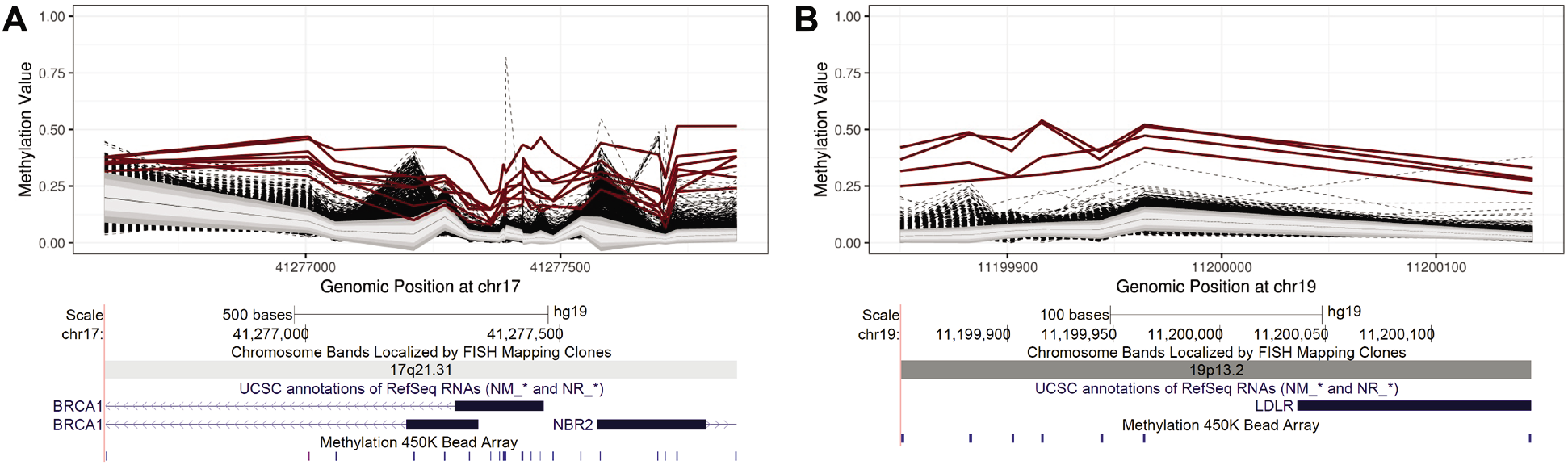
Multiple individuals with epivariations at the promoters of *BRCA1* and *LDLR*. Coding mutations of *BRCA1* and *LDLR* are associated with familial breast/ovarian cancer, and familial hypercholesterolemia, respectively. Loss of function mutations in both genes are considered highly penetrant and medically actionable, and are recommended for return to patients when identified as incidental or secondary findings in clinical sequencing under current guidelines published by the American College of Medical Genetics and Genomics^39^. We identified promoter hypermethylation of *BRCA1* in seven individuals, and promoter hypermethylation of *LDLR* in four individuals, yielding population estimates for these epivariations of approximately 1 per 3,300 and 1 per 5,800 individuals, respectively. In each methylation plot, individuals with epivariations are shown as bold red lines, while the other ~23,000 samples tested are shown as dashed black lines. Grey shaded regions indicate 1, 1.5 and 2 Standard Deviations from the mean of the distribution, while the solid black line shows the mean β-value of the entire population. Below each methylation plot are screenshots from the UCSC Genome Browser showing the relevant genomic region, genes, and the location of CpGs assayed by probes on the Illumina 450k array.

### Segregation of epivariations with local sequence variants

We hypothesized that some epivariations might represent secondary events that segregate within the population on consistent haplotype backgrounds^7,9^. Using data from 933 individuals from the Women’s Health Initiative in whom both methylation and single nucleotide variation (SNV) data were available, we identified 97 epivariations that were present in at least two individuals, and performed association analysis of these with local SNVs. Overall, using a stringent statistical threshold (1% FDR), 68 epivariations (70%) showed at least one significantly associated SNV (Figure 4, Table S8). There was a trend for significantly associated variants to be located in close proximity to the epivariation, and the region of significantly associated SNVs directly overlapped the epivariation in many cases. These results indicate that many epivariations result from genetic variants located within their immediate vicinity^10^. However, in a few cases, significant associations occurred with SNVs located >500 kb away, suggesting that some genetic variants can disrupt epigenetic regulation over large distances *in cis* (Figure S3).

**Figure 4.**
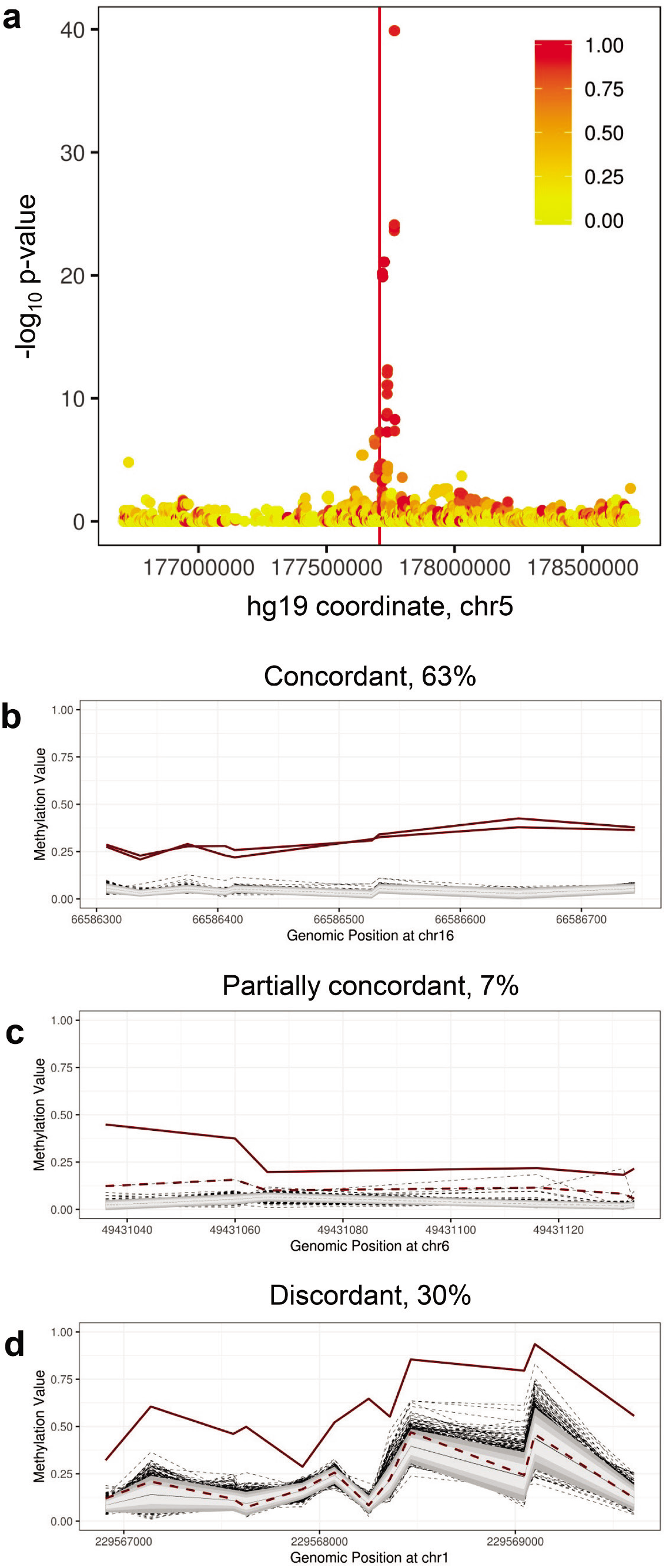
Genetic association and twin concordance analysis provides insight into the origins of epivariations. **(A)** Many epivariations are associated with local sequence variation, indicating an underlying genetic cause. Using data from 933 individuals from the Women’s Health Initiative in whom both methylation and SNV data were available, we performed association analysis with local SNVs for each recurrent epivariation. Shown are results for an epivariation located intronic within *COL23A1* [MIM: 610043] (chr5:177,707,036-177,707,227, hg19) that showed a strong association with local SNVs. Eight of the nine epivariation carriers were heterozygous for a cluster of 12 SNVs (lead SNV p=1.3×10^−40^, χ-square test), each of which had minor allele frequency of only 1.3% in non-carriers. The region of significant association directly overlaps the epivariation, indicating that this epivariation is likely a secondary event that occurs as a result of genetic variation segregating on a local haplotype. The color of each point indicates the fraction of epivariation carriers that carry the associated allele, while the location of the epivariation is indicated by the vertical red line. **(B-D)** One third of epivariations are discordant in monozygotic twin pairs, suggesting a high frequency of somatic epivariation. We identified 333 epivariations in 700 pairs of MZ twins. Of these, 63% were concordant with both members of the twin pair carrying the same epivariation, 7% were partially concordant, with one twin carrying the epivariation and the second twin being a weak outlier, and 30% were discordant with only one of the two twins carrying the epivariation. **(B)** chr16:66,586,308-66,586,746, located at the promoter of *CKLF* [MIM: 616074]. **(C)** chr6:49,431,037-49,431,136, located at the bidirectional promoter of *CENPQ* [MIM: 611506] and *MMUT* [MIM: 609058], the latter of which is associated with methylmalonic aciduria. **(D)** chr1:229,566,907-229,569,609, overlapping *ACTA1* [MIM: 102610], which is associated with a number of congenital myopathies. In each methylation plot, the two members of an MZ twin pair are shown as bold red lines. Individuals formally identified as carrying an epivariation by our sliding window analysis are shown as solid red lines, while a member of the twin pair who was not formally identified as carrying this epivariation are shown as dashed red lines. All other samples from the cohort are shown as dashed black lines, with the grey shaded regions indicating 1, 1.5 and 2 Standard Deviations from the mean of the distribution, while the solid black line shows the mean β-value of the entire population.

### Epivariations are frequently discordant in monozygous twins

As MZ twins arise from the splitting of a single embryo post-fertilization, they provide a unique opportunity to gain insights into the developmental origins of epivariations. Our study population included 700 pairs of MZ twins derived from four different cohorts, and we identified a total of 333 loci where epivariations were identified in one or both of these MZ twins. Manual curation of these events showed that while 63% were concordant (*i.e*., both members of the MZ twin pair carried the same epivariation), 30% showed complete discordance (where one twin carried the epivariation, while the second twin showed a normal methylation pattern at that locus), and 7% were scored as partially concordant (where one twin carried the epivariation, while the second twin showed an outlier methylation profile at that locus, but of reduced magnitude) (Table S9). Examples of these three categories are shown in Figure 4. Overall, these observations indicate that approximately one third of epivariations are somatic events that occur post-zygotically.

### Epivariations are more common with age

Using 11,690 samples with reported age at sampling, we observed a significant trend for the number of epivariations identified per individual to increase with age (Spearman r=0.17, p=4×10^−81^) (Figure 5). Consistent with this, MZ twin pairs who were fully or partially discordant for an epivariation (mean age 36.5 years) were significantly older than MZ twins with concordant epivariations (mean age 26.2 years) (p=0.002, two-sided t-test). These observations suggest that some epivariations are sporadic somatic events that accumulate with age.

**Figure 5.**
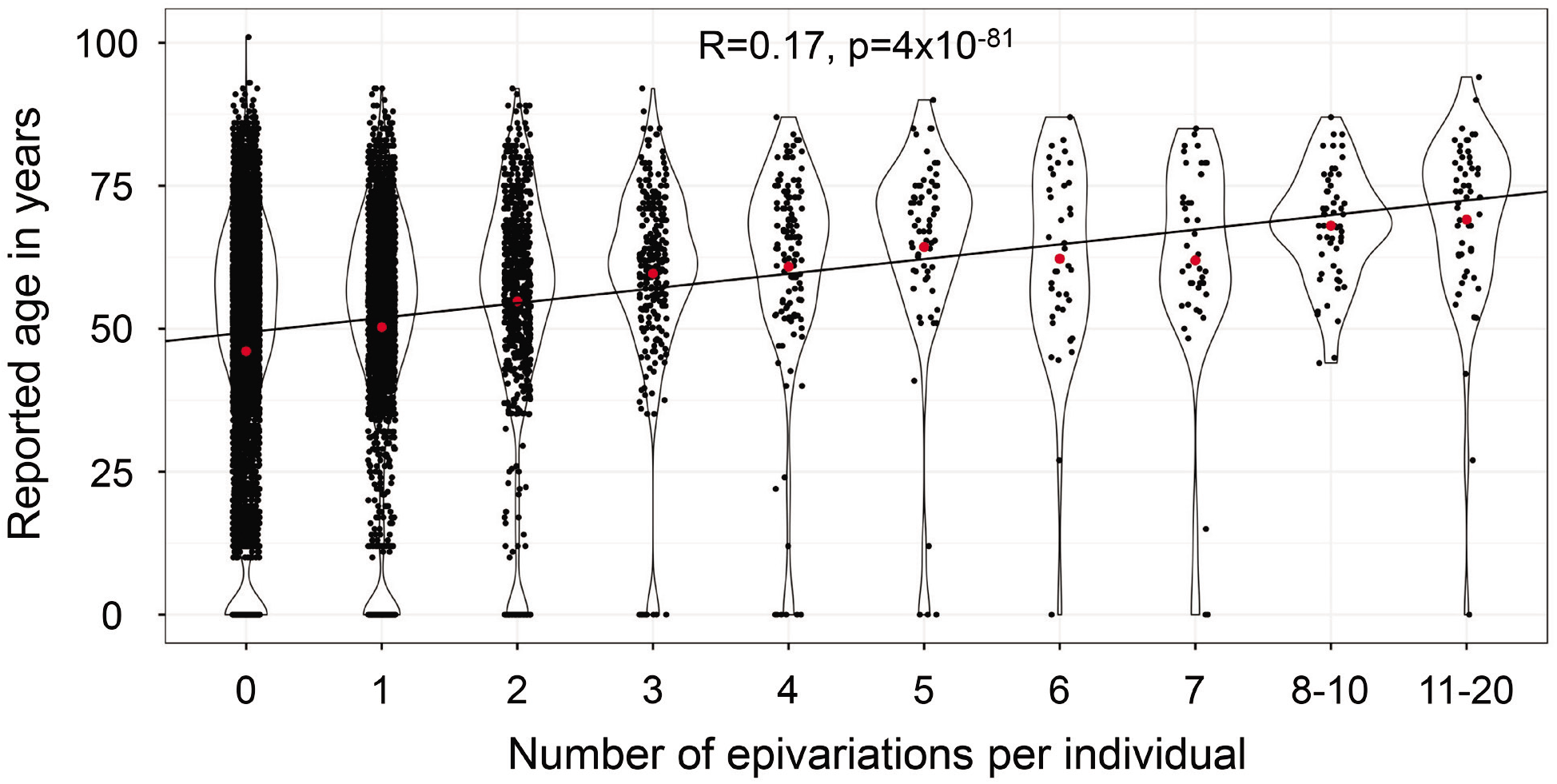
Epivariations are more common with age. Using 11,690 individuals with reported age at sampling and methylation profiled in blood, we observed a significant trend for the number of epivariations per individual to increase with age (Spearman r=0.17, p=4×10^−81^). The red dot in each violin shows the group mean.

### Epivariations at imprinted loci

Epivariations at imprinted loci exhibited a frequency profile that differed from the overall genomic distribution of epivariations in several ways: (i) Imprinted loci were more prone to epivariations, showing a 4.2-fold increase in epivariations compared to non-imprinted loci (p=8.5×10^−22^, Hypergeometric test). (ii) Loss of methylation defects predominate over gains of methylation at imprinted loci: 60% of epivariations at imprinted loci were hypomethylation events, representing a 2.3-fold increase compared to the rest of the genome (p=6.3×10^−45^, Hypergeometric test) (Figure S4). (iii) Consistent with their hemi-methylated nature, epivariations at imprinted loci were 85-fold enriched for bi-directional changes compared to the entire genome (p=1.2×10^−24^, Hypergeometric test) (Figure S5).

### Prediction and validation of novel CGG expansions at hypermethylated epivariations

Using tandem repeat (TR) genotypes generated by *hipSTR* in 600 unrelated individuals who had undergone Illumina WGS, we observed that TRs that are known to undergo occasional expansion in human disease tend to show extremely high levels of polymorphism in the general population (Figure S6). For example, nearly all known pathogenic TRs had ≥10 different alleles in the 600 genotyped samples, placing them in the top 3% of the most polymorphic TRs in the genome. Thus, we hypothesized that we could use high levels of population variability to predict unstable TRs in the human genome that are prone to occasional expansion. Based on this approach, we identified 180 TRs with motif size of 3-6 bp and 100% GC-content that each showed ≥10 different alleles in our cohort of 600 sequenced individuals. Intersection of these potentially unstable GC-rich TRs with hypermethylated epivariations yielded 25 overlaps. This included six TRs that were already known to undergo rare expansion and hypermethylation (*FRA10AC1, C11orf80* [MIM: 616109], *CBL* [MIM: 165360], *C9orf72* [MIM: 614260], *DIP2B*, and *TMEM185A* [MIM: 300031])^15,18,19,37,40–42^, and highlighted 19 additional epivariations that we hypothesized might be caused by previously unidentified TR expansions (Table S10). Plots of all methylation profiles of epivariations overlapping putatively unstable CG-rich TRs are shown in Figure S7.

To investigate whether these epivariations were attributable to expansions of an underlying TR, we obtained DNA samples from four individuals in whom we had identified hypermethylated epivariations overlapping putatively unstable CGG repeats and performed long-read WGS using either Pacific Biosciences SMRT sequencing or Oxford Nanopore Technology (ONT). In all four samples tested, we confirmed the presence of a heterozygous TR expansion comprising several hundred copies of CGG at the epivariation (Figure 6, Table S10), thus identifying novel hypermethylated CGG expansions within the promoter/5’UTR regions of *ABCD3* [MIM: 170995], *FZD6/LOC105369147* [MIM: 603409], *LINGO3* [MIM: 609792] and *PCMTD2*. Furthermore, by analyzing the signal profiles of phased ONT reads, we demonstrated that in an individual with hypermethylation of *FZD6*, the expanded TR allele was highly methylated, while the normal TR allele was largely unmethylated (Figure 6b), thus showing that this epivariation represents monoallelic hypermethylation associated with a CGG expansion.

**Figure 6.**
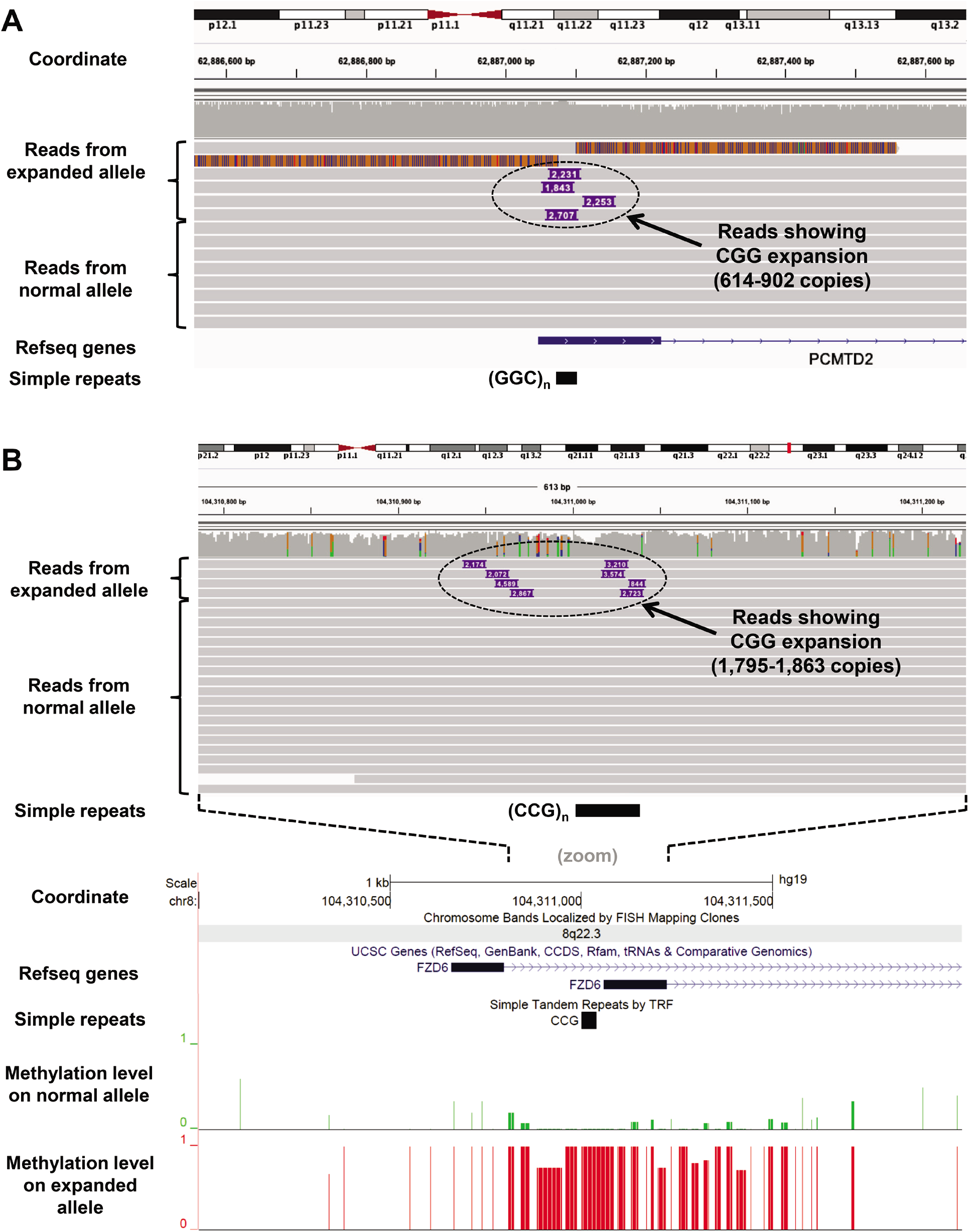
Validation of novel CGG expansions underlying epivariations using long read sequencing. We obtained DNA samples from individuals with hypermethylated epivariations at the promoter/5’ UTR regions of *PCMTD2* and *FZD6*, both of which contained putatively unstable CGG repeats. **(A)** Pacific Biosciences SMRT sequencing confirmed the presence of a heterozygous repeat expansion at *PCMTD2* composed of 614-902 CGG motifs, suggesting considerable somatic variation in the size of the expanded allele. **(B)** Sequencing with Oxford Nanopore Technology confirmed the presence of a heterozygous repeat expansion at *FZD6*, composed of 1,795-1,863 CGG motifs. By analyzing the signal profiles of the reads after phasing based on presence/absence of the expansion, we demonstrated that the expanded allele was highly methylated, while the normal TR allele was largely unmethylated.

Multiple folate-sensitive fragile sites (FSFS) in the human genome are known to be caused by underlying CGG expansions, including FRA2A (*AFF3*), FRA7A (*ZNF713*, [MIM: 616181]), FRA10A (*FRA10AC1*), FRA11A (*C11orf80*), FRA11B (*CBL*), FRA12A (*DIP2B*), FRA16A (*XYLT1*), FRAXA (*FMR1*), FRAXE (*AFF2*), and FRAXF (*TMEM185A*)^16^,^17^,^45^,^18–20,37,40,41,43,44^. We thus hypothesized that novel CGG expansions might underlie other FSFS. Consistent with this, eight of the 25 putative and validated CGG repeat expansions we identified coincide with the cytogenetic location of other rare FSFS that to date have not been molecularly mapped^16^, strongly suggesting that these epivariations likely represent the unstable CGG repeats that are responsible for the FSFS FRA1M (*ABCD3*), FRA2B (*BCL2L11*, [MIM: 616181]), FRA5G (*FAM193B*, [MIM: 615813]), FRA8A (*FZD6*), FRA9A (*C9orf72*) FRA19B (*LINGO3*), FRA20A (*RALGAPA2*) and FRA22A (*CSNK1E*, [MIM: 600863]) (Table S10).

To formally test whether this approach accurately identifies CGG expansions underlying FSFS, we obtained DNA from an individual who expressed the FSFS FRA22A, but who was not part of our discovery cohort. Our epivariation analysis had identified six individuals with a gain of methylation overlapping the 5’UTR of *CSNK1E*, a region that includes a highly polymorphic CGG repeat and lies within 22q13.1, the cytogenetic band to which FRA22A has been mapped. Thus, based on our epivariation and TR data, we predicted that expansions of this CGG repeat within *CSNK1E* likely underlie the FRA22A fragile site, which was subsequently confirmed by several complementary experiments: (i) Repeat primed-PCR of the CGG repeat^46^ in the individual with the FRA22A fragile site showed a characteristic saw-tooth pattern on the fluorescence trace, with periodicity of 3 bp, indicative of a triplet repeat expansion. (ii) Subsequent Southern blot in the FRA22A carrier identified a novel smeared fragment of approximately 3.2-kb, in addition to the expected fragment of 2.2-kb, which, together with the PCR result, indicate the presence of an expanded CGG tract of approximately 340 repeats. (iii) Analysis of CpGs in the promoter of *CSNK1E* using both bisulfite sequencing and pyrosequencing showed methylation levels of 40-50% in the FRA22A carrier, while control samples were unmethylated. (iv) Finally, using real-time RT-PCR in lymphoblastoid cells, we observed that in the FRA22A carrier, expression of *CSNK1E* was reduced to ~37% of the levels observed in controls (Figure S8). Overall, these results indicated that expansion of a CGG repeat in the 5’UTR of *CSNK1E* results in allelic methylation and silencing of the gene, and represents the molecular defect underlying the FRA22A FSFS.

## DISCUSSION

Our large-scale survey of epivariations in >23,000 individuals represents the largest cohort of methylomes assembled to date, providing a comprehensive catalog of epivariations that are found in the human population. While a handful of previous studies have identified epivariations as causative factors in some human genetic diseases, here we identified promoter epivariations at hundreds of genes that are known to cause genetic diseases, suggesting that epivariations may contribute to the mutational spectra underlying many Mendelian diseases. Using available expression data, we show that many of these epivariations exert functional effects on the genome, with promoter epivariations in particular being associated with significant alterations in gene expression. In previous work, we have shown that hypermethylated promoter epivariations are often associated with monoallelic expression, and thus can have an impact comparable to that of loss-of-function coding mutations^10^. Based on this observation, we anticipate that epigenetic profiling in patients with overt genetic disease, but who lack pathogenic sequence mutations in the gene(s) relevant to their phenotype, will lead to the identification of epivariants as a causative factor in some conditions, and potentially providing additional diagnostic yield compared to purely sequence based approaches^11^.

Through genetic association and by studying patterns of epivariation in twin pairs and samples of different ages, we gained insights into the underlying mechanisms of epivariations. These analyses suggest that ~70% of epivariations segregate on specific haplotype backgrounds, indicating the majority of epivariations are secondary events that occur downstream of stably inherited genetic variants. We hypothesize that these epivariations result from underlying sequence variants that disrupt either the establishment of maintenance of the normal epigenetic state, such as mutations of regulatory elements and transcription factor binding sites^10,47^. Contrastingly, analysis of MZ twins found that approximately one third of epivariations are discordant between genetically identical twins, indicating that a significant fraction of epivariations occur post-zygotically. This conclusion is further supported by the observations that (i) the incidence of epivariations increases with age, and (ii) in MZ twins, discordant epivariations are observed more frequently in older versus younger twins. This suggests that many epivariations will likely exhibit somatic mosaicism and therefore, depending on their tissue distribution, might show attenuated or absent phenotypic effects, and reduced heritability between generations. Consistent with this latter prediction, we previously observed a significant reduction in heritability of epivariations between parents and offspring^10^. We postulate that post-zygotic epivariations may represent either (i) primary epivariations, *i.e*., sporadic errors that arise as a result of the epigenetic remodelling that occurs during cellular differentiation, or (ii) secondary epivariations resulting from somatically-acquired sequence mutations. Further work will be needed to distinguish these possibilities.

We also identified CGG repeat expansions as the causative factor underlying a subset of epivariations. Large expansions of CGG repeats are known to be associated with local DNA hypermethylation of the expanded allele, and have been found to underlie multiple rare folate-sensitive fragile sites in the genome^16^. By combining our map of outlier hypermethylation events with predictions of unstable TRs, we identified 25 epivariations that we predicted as being caused by underlying CGG repeat expansions. Six of these loci represent previously identified TR expansions, thus both validating our approach, and providing population estimates of the prevalence of these events, some of which are surprisingly frequent. For example, our data indicate that hypermethylated expansions at *FRA10AC1* occur with a prevalence of ~1 per 330 individuals. In order to assess the validity of our predictions for the nineteen other loci containing CGG repeats, we obtained DNA samples from five individuals in whom we identified hypermethylation of the candidate loci, and validated the presence of a heterozygous expanded repeat at all five of these loci in the carrier individuals. Although we were unable to obtain DNA samples with putative expansions at the 14 other putatively unstable CGG TRs we identify, we suggest that these represent strong candidates for novel TR expansions. In support of this, several of these candidate loci coincide with the approximate location of rare FSFSs that have been cytogenetically mapped, suggesting that these candidate repeats represent the molecular defect underlying these FSFSs. While several hypermethylated CGG expansions are known to be associated with neurodevelopmental disorders^17–20,43,45^, the possible phenotypic consequences of the novel CGG expansions we identified will require further study. Given that many of these occur within the 5’ UTRs of genes, one intriguing possibility is that unmethylated premutation-sized alleles might predispose to late-onset neurodegenerative disease, similar to the Fragile X tremor/ataxia syndrome that occurs in some carriers of *FMR1* premutations^48^.

In an era where genome sequencing is being applied to millions of individuals, our results show that the study of epigenetic variation can provide additional insights into genome function.

## DESCRIPTION OF SUPPLEMENTAL DATA

Supplemental Data include eight Supplemental Figures, and ten Supplemental Tables.

## ACKNOWLEDGEMENTS

This work was supported by NIH grant NS105781 to AJS, NIH predoctoral fellowship NS108797 to OR, and American Heart Foundation Postdoctoral Fellowship 18POST34080396 to AMT. R.F.K acknowledges support of the Research Fund of the University of Antwerp (Methusalem-OEC grant – “GENOMED”). Research reported in this paper was supported by the Office of Research Infrastructure of the National Institutes of Health under award number S10OD018522. The content is solely the responsibility of the authors and does not necessarily represent the official views of the National Institutes of Health. This work was supported in part through the computational resources and staff expertise provided by Scientific Computing at the Icahn School of Medicine at Mount Sinai.

The Biobank-Based Integrative Omics Studies (BIOS) Consortium is funded by BBMRI-NL, a research infrastructure financed by the Dutch government (NWO 184.021.007). The Parkinson’s disease patient and control study was funded by NIEHS grants ES024356, R01ES10544, and P01ES016732. The Framingham Heart Study is conducted and supported by the National Heart, Lung, and Blood Institute (NHLBI) in collaboration with Boston University (Contract No. N01-HC-25195 and HHSN268201500001I). Additional funding for SABRe was provided by Division of Intramural Research, NHLBI, and Center for Population Studies, NHLBI. The Women’s Health Initiative (WHI) program is funded by the National Heart, Lung, and Blood Institute, National Institutes of Health, U.S. Department of Health and Human Services through contracts HHSN268201600018C, HHSN268201600001C, HHSN268201600002C, HHSN268201600003C, and HHSN268201600004C. This manuscript was not prepared in collaboration with investigators of the Framingham Heart Study or WHI, and does not necessarily reflect the opinions or views of the Framingham Heart Study, WHI investigators, Boston University, or NHLBI.

## WEB RESOURCES

Array Express, https://www.ebi.ac.uk/arrayexpress/

Beagle, http://bochet.gcc.biostat.washington.edu/beagle/1000_Genomes_phase3_v5

Database of Genotypes and Phenotypes (dbGaP), https://www.ncbi.nlm.nih.gov/gap/

European Genome-phenome Archive, https://www.ebi.ac.uk/ega/home

Gene Expression Omnibus (GEO), https://www.ncbi.nlm.nih.gov/geo/

OMIM, http://www.omim.org/

UCSC Genome Browser, http://genome.ucsc.edu

All code used in this paper are available at https://github.com/AndyMSSMLab/Epivariation-in-23K-samples

